# *Vibrio tarriae* sp. nov., a novel member of the Cholerae clade isolated from across the United States

**DOI:** 10.1101/2022.04.17.488585

**Authors:** Mohammad Tarequl Islam, Kevin Liang, Fabini D. Orata, Monica S. Im, Munirul Alam, Christine C. Lee, Yann F. Boucher

## Abstract

A number of bacteria with close resemblance to *Vibrio cholerae* has been isolated over the years by the Centers for Disease Control and Prevention (CDC), which could not be assigned a proper taxonomic designation based on preliminary identification methods. Nine such isolates have been found to share 16S rRNA gene identity exceeding 99% with *V. cholerae*, yet DNA-DNA hybridization (60.4-62.1%) and average nucleotide identity values (94.4-95.1%) were below the species cut-off, indicating a potentially novel species. Phylogenetic analysis of core genomes places this group of isolates in a monophyletic clade, within the “Cholerae clade,” but distinct from any other species. Extensive phenotypic characterization reveals unique biochemical properties that distinguish this novel species from *V. cholerae*. Comparative genomic analysis reveals a unique set of siderophore genes, suggesting that iron acquisition strategies could be vital for the divergence of the novel species from a common ancestor with *V. cholerae*. Based on genetic, phylogenetic, and phenotypic differences observed, we propose these isolates represent a novel species of the genus *Vibrio*, for which the name *Vibrio tarriae* sp. nov. is proposed. Strain 2521-89 (= DSM 112461 = CCUG 75318), isolated from lake water, is the type strain.

**Author Notes:** The GenBank/EMBL/DDBJ accession number for the 16S rRNA gene sequence of strain 2521-89 is MW773202.1. The genome sequences (genome assemblies) of strains 2521-89, 2523-88, 2015V-1076, 2016V-1018, 2016V-1062, 2017V-1038, 2017V-1070 and 2017V-1124 are deposited under the accession number NZ_CP022353.1, QKKG01000001.1, QKKH00000000.1, QKKI00000000.1, QKKJ00000000.1, QKKK00000000.1, QKKM00000000.1, and QKKN00000000.1, respectively. All the whole genome sequences are deposited under bioproject ID: PRJNA391152.

## Introduction

‘Vibrios’ are globally important organisms, frequently causing diseases in humans and aquatic animals as well as serving as potential indicators of climate change [1, 2]. From a clinical perspective, the Cholerae clade represents one of the most important groups of bacteria among the diverse assemblages within the genus *Vibrio* [3]. The clade is termed after *V. cholerae*, the type strain of the genus and the causative agent of cholera, a devastating pandemic disease [4]. *V. cholerae* is an extensively studied bacteria, largely because of the lethality of certain lineages. It is also a model system for environmental pathogens causing human disease [5]. Despite this, our understanding of the natural diversity of *V. cholerae* and its close relatives have been limited until recently [6, 7]. As a result of some notable environmental surveillance efforts, a few closely related species have recently been found; most were initially classified as *V. cholerae*-like bacteria in the last few decades [6, 8–10]. In the U.S., *Vibrio*-related diseases are on the rise, affecting an increasing number of people coming in contact with water during recreational or professional activities [11]. Global climate change is thought to have contributed to this alarming increase in cases [11, 12] as vibrios are aquatic bacteria and their ecology is tightly correlated with environmental conditions [2, 13]. To track the emergence of novel and potentially dangerous species, surveillance is of paramount importance. The National *Vibrio* Reference Laboratory at the Centers for Disease Control and Prevention (CDC) have collected and identified potential *Vibrio* pathogens from human clinical specimens across the U.S. while national surveillance is conducted under the Cholera and Other *Vibrio* Illness Surveillance (COVIS) program [14].

While the pathogenic potential of species related to *V. cholerae* vary and severe illness is rare, cases associated with opportunistic infections can be lead to serious complications [11]. Furthermore, the interaction of *V. cholerae* with other pathogenic bacteria in nature and inside the human gut can lead to the emergence of distinct pathogens. For example, closely related species have been found to exchange genetic material with *V. cholerae*, including genes encoding virulence factors. For example, *Vibrio mimicus* contains virulence genes including the cholera toxin and was initially isolated from diarrheal patients [10]. In the environment, *Vibrio metoecus* frequently exchanges genes via horizontal gene transfer (HGT) with *V. cholerae* [15]. The recently described closest known sister species of *V. cholerae, Vibrio paracholerae*, had historically been associated with human infections and co-exists with *V. cholerae* in cholera endemic areas [6, 16].

In this study, we employed a polyphasic approach including phenotypic, genetic, and phylogenetic data to describe a novel species isolated from both human and environmental samples from across the U.S. The name *V. tarriae* sp. nov. is proposed for this novel species to honor the decades of public service of Dr. Cheryl L. Tarr in fighting vibriosis and cholera across the world and her contribution to our understanding of *Vibrio* taxonomy, physiology, and epidemiology.

## Materials and Methods

### Bacterial isolates

The bacterial isolates were collected by the CDC as part of the national reference laboratories and includes isolates from clinical cases across the U.S. Nine isolates were identified as *Vibrio* spp. by preliminary culture identification (Table 1); genome sequencing and preliminary genomic analyses identified the strains as a close relative of *V. cholerae* [17, 18]. The isolates were transported to the University of Alberta following standard bacterial isolate transportation procedures [19] for further analysis.

**Table 1.**
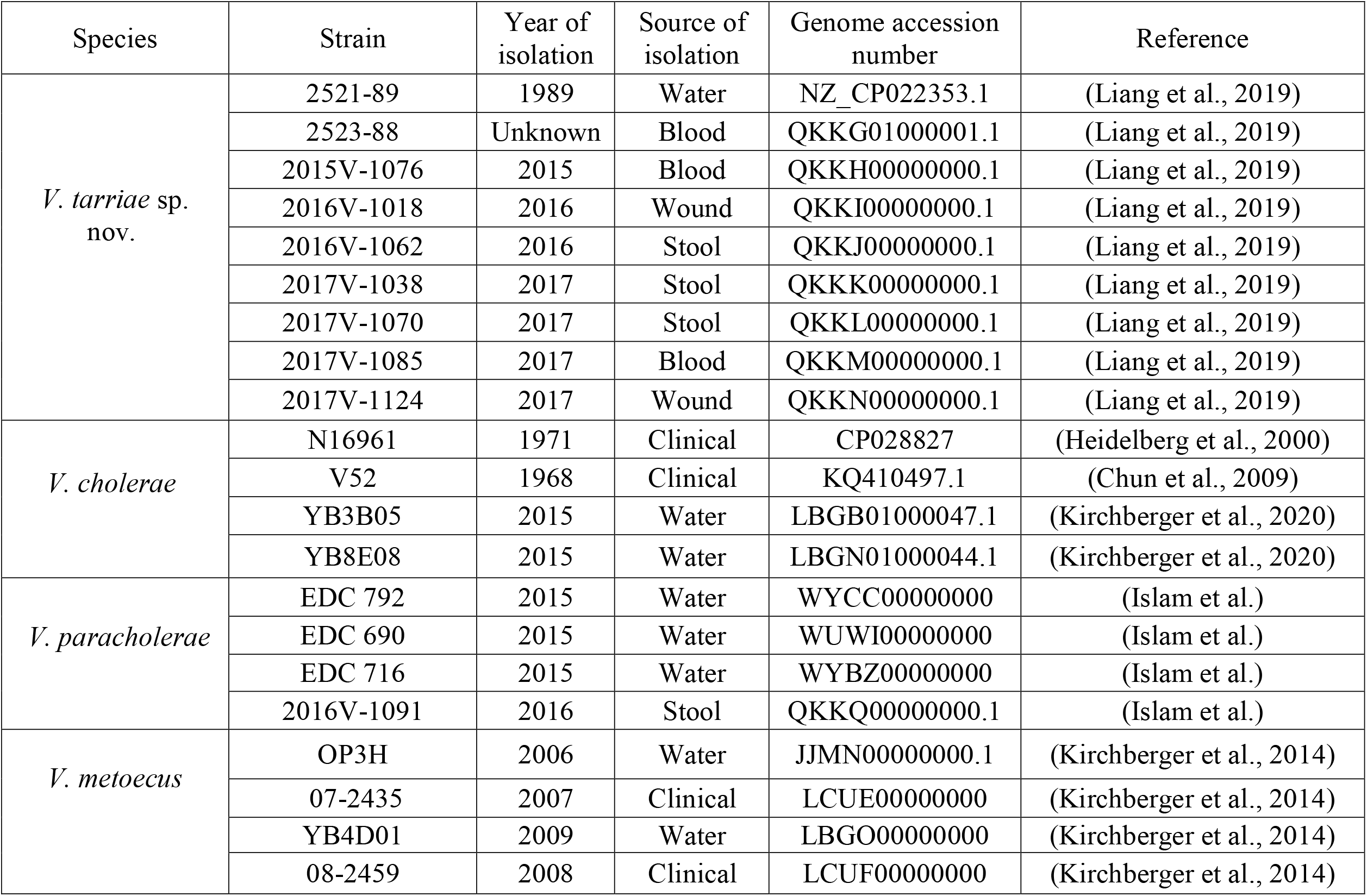
Information of *Vibrio tarriae* sp. nov. strains along with the reference strains of closely related species: *V. cholerae, V. paracholerae* and *V. metoecus* used in this studys.

| Species | Strain | Year of isolation | Source of isolation | Genome accession number | Reference |
| --- | --- | --- | --- | --- | --- |
| *V. tarriae* sp. nov. | 2521-89 | 1989 | Water | NZ_CP022353.1 | (Liang et al., 2019) |
|  | 2523-88 | Unknown | Blood | QKKG01000001.1 | (Liang et al., 2019) |
|  | 2015V-1076 | 2015 | Blood | QKKH00000000.1 | (Liang et al., 2019) |
|  | 2016V-1018 | 2016 | Wound | QKKI00000000.1 | (Liang et al., 2019) |
|  | 2016V-1062 | 2016 | Stool | QKKJ00000000.1 | (Liang et al., 2019) |
|  | 2017V-1038 | 2017 | Stool | QKKK00000000.1 | (Liang et al., 2019) |
|  | 2017V-1070 | 2017 | Stool | QKKL00000000.1 | (Liang et al., 2019) |
|  | 2017V-1085 | 2017 | Blood | QKKM00000000.1 | (Liang et al., 2019) |
|  | 2017V-1124 | 2017 | Wound | QKKN00000000.1 | (Liang et al., 2019) |
| *V. cholerae* | N16961 | 1971 | Clinical | CP028827 | (Heidelberg et al., 2000) |
|  | V52 | 1968 | Clinical | KQ410497.1 | (Chun et al., 2009) |
|  | YB3B05 | 2015 | Water | LBGB01000047.1 | (Kirchberger et al., 2020) |
|  | YB8E08 | 2015 | Water | LBGN01000044.1 | (Kirchberger et al., 2020) |
| *V. paracholerae* | EDC 792 | 2015 | Water | WYCC00000000 | (Islam et al.) |
|  | EDC 690 | 2015 | Water | WUWI00000000 | (Islam et al.) |
|  | EDC 716 | 2015 | Water | WYBZ00000000 | (Islam et al.) |
|  | 2016V-1091 | 2016 | Stool | QKKQ00000000.1 | (Islam et al.) |
| *V. metoecus* | OP3H | 2006 | Water | JJMN00000000.1 | (Kirchberger et al., 2014) |
|  | 07-2435 | 2007 | Clinical | LCUE00000000 | (Kirchberger et al., 2014) |
|  | YB4D01 | 2009 | Water | LBGO00000000 | (Kirchberger et al., 2014) |
|  | 08-2459 | 2008 | Clinical | LCUF00000000 | (Kirchberger et al., 2014) |

### Phenotypic tests

The isolates were sub-cultured on thiosulfate citrate bile salts sucrose (TCBS) agar (Becton Dickinson) and TSB agar (Difco) and incubated overnight at 30°C. Single colonies from the TSB agar cultures were tested using the Analytical Profile Index (API) 20 NE (bioMerieux), the Phenotype MicroArray 1 (PM1) and 2 (PM2) microplate (Biolog) according to manufacturer’s instructions. Gram staining and viewing of the cells under a light microscope (Carl Zeiss) at 1000X magnification was performed [20]. Permissive growth temperatures were determined in Brain heart infusion broth (BHI) with incubation at a range of 4–45□, whereas permissive salinity concentrations were determined in TSB at 30□ in a range of 0–10 % NaCl.

### Genetic and bioinformatic analysis

For genetic characterization, reference genome sequences were obtained from the GenBank database and accession numbers are listed in Table 1. The G+C content was determined from whole-genome sequences using Geneious 8.1.2 [21]. Pairwise average nucleotide identity (ANI) was calculated using JSpecies V1.2.1 [22]. Pairwise percent DNA–DNA hybridization (dDDH) was also calculated *in silico* using the Genome-to-Genome Distance Calculator 2.0 (GGDC) [23]. The genome sequences were annotated with RAST 2.0 [24]. Core and accessory genes were determined with BPGA finding orthologous protein-coding genes clustered into families based on a 30% amino acid sequence identity [25]. The gene families unique to any particular group were determined using a custom Python program. Genome sequences were aligned using Mugsy [26] and the alignments were concatenated, stripping columns with at least one gap, using Geneious 8.1.2. This resulted i n a single alignment with a total length of 972,240 bp, which was used to reconstruct a maximum-likelihood tree with RAxML 8.2.8 [27] using the GTR (general time reversible) nucleotide substitution model with gamma distribution of rate categories and 100 bootstrap replicates. For multi-locus sequence analysis (MLSA), six housekeeping genes *adk, gyrB, pyrH, pgi, recA* and *rpoA* were selected [8, 20, 28]. From the DNA sequences, a concatenated alignment of 7,392 bp was obtained and used to reconstruct a maximum-likelihood tree.

## Results and Discussion

### Distinctive phenotypic traits of the novel species

Phenotypic characterization was performed on four representative isolates of *V. tarriae* sp. nov. and compared with same number of isolates from *V. cholerae, V. paracholerae* and *V. metoecus* (Tables 1). Overall phenotypic traits support inclusion in the cholerae clade. All the *V. tarriae* sp. nov. isolates studied exhibited growth in TSB without NaCl. Ability to grow in the media without additional salt is characteristic of the clade including *V. cholerae, V. paracholerae, V. metoecus* and *V. mimicus*, differentiating it from the rest of the vibrios [3, 29]. The isolation of strain 2521-89 of *V. tarriae* sp. nov. from lake water [17] suggests that the species is able to survive in freshwater environments. Furthermore, the isolates were also able to survive at 40 °C, consistent with other pathogenic vibrios that can survive inside the human body [29].

*Vibrio tarriae* sp. nov. resembles *V. cholerae* (160 of 190 tests or 84%; Table 2) and *V. paracholerae* (172 of 190 tests or 90%) in the majority of phenotypic characteristics tested. However, eleven phenotypic features distinguished *V. tarriae* sp. nov. from *V. cholerae* and at least two from *V. paracholerae* (summarized in Table 2). Notably, the new species tested positive for the utilization of N-acetyl galactose amine and pectin as the sole carbon and energy source, while *V. cholerae* was negative for these tests. In contrast to *V. cholerae*, these strains were unable to utilize D-mannose, citric acid, propionic acid, monomethyl succinate, caproic acid, D-Serine, D-Glucose-6-phosphate, glycyl-L-proline and D-fructose-6-phosphate. It also differs from *V. paracholerae* by two phenotypic traits: the ability to utilize N-acetyl galactose amine and inability to utilize α-cyclodextrin. Species of the cholerae clade are usually known to be lactose negative, and the ability to ferment lactose is a distinguishing characteristic for *V. vulnificus* [30]. In our phenotypic tests, all four *V. tarriae* sp. nov. isolates and one *V. paracholerae* isolate were positive for growth on lactose, whereas all *V. cholerae* and *V. metoecus* isolates were negative for lactose utilization (Table 2).

**Table 2:** Summary of phenotypic test results for *V. tarriae* sp. nov., *V. cholerae* and *V. paracholerae* and *V. metoecus* strains: 1, *V. tarriae* sp. nov. 2521-89; 2, V. *tarriae* sp. nov. 2015V-1076; 3, V. *tarriae* sp. nov. 2017V-1070; 4, V. *tarriae* sp. nov. 2017V-1038; 5, *V. cholerae* N16961; 6, *V. cholerae* V52; 7, *V. cholerae* YB3B05; 8, *V. cholerae* YB8E08; 9, *V. paracholerae* EDC 690; 10, *V. paracholerae* EDC 792; 11, *V. paracholerae* 2016V-1091; 12, *V. paracholerae* 2016V-1114; 13, *V. metoecus* OP3H; 14. *V. metoecus* OP4B; 15, *V. metoecus* OP6B; *V. metoecus* Vm082459; +, Growth/positive test result; -, no growth/negative test result.

| Phenotypic test | *Vibrio tarriae* sp. nov. |  |  |  | *Vibrio cholerae* |  |  |  | *Vibrio paracholerae* |  |  |  | *Vibrio metoecus* |  |  |  |
| --- | --- | --- | --- | --- | --- | --- | --- | --- | --- | --- | --- | --- | --- | --- | --- | --- |
|  | 1 | 2 | 3 | 4 | 5 | 6 | 7 | 8 | 9 | 10 | 11 | 12 | 13 | 14 | 15 | 16 |
| D-Lactose | + | + | + | + | - | - | - | - | - | - | - | + | - | - | - | - |
| N-Acetyl-D-Galactosamine | + | + | + | + | - | - | - | - | - | - | - | - | + | + | + | + |
| Pectin | + | + | + | + | - | - | - | - | + | + | + | + | + | + | + | + |
| D-Mannose | - | - | - | - | + | + | + | + | - | - | - | - | + | + | + | + |
| D-Serine | - | - | - | - | + | + | + | + | - | - | - | + | + | + | + | + |
| Citric Acid | - | - | - | - | + | + | + | + | - | - | - | - | - | - | - | - |
| D-Ribose | - | - | - | - | + | + | + | + | - | - | - | - | - | - | - | - |
| Propionic Acid | - | - | - | - | + | + | + | + | - | - | - | - | - | - | - | - |
| D-Alanin | - | - | - | - | + | + | + | + | - | - | + | - | - | - | - | - |
| Mono Methyl Succinate | - | - | - | - | + | + | + | + | - | - | - | - | - | - | - | - |
| Caproic Acid | - | - | - | - | + | + | + | + | - | - | - | - | - | - | - | - |
| α-cyclodextrin | - | - | - | - | - | - | - | - | + | + | + | + | - | + | + | + |

### Genome relatedness and phylogenetic position in respect to well-known relatives

The DNA G+C content of the *V. tarriae* sp. nov. isolates ranged from 47.1–47.2%, which is consistent with the known range for the genus *Vibrio* (38.0–51.0%) [29]. The 16S rRNA gene sequences were >99% identical to *V. cholerae, V. paracholerae, V. metoecus*, and *V. mimicus* strains in the NCBI database. DNA-DNA relatedness of *V. tarriae* sp. nov strains with these species were determined using two matrixes: average nucleotide identity (ANI) and digital DNA-DNA hybridization (dDDH) by pairwise comparisons of whole-genome sequences. The ANI values between isolates within *V. tarriae* sp. nov. range from 98-99% (Fig. 1). In contrast, the ANI values for *V. tarriae* sp. nov. was 94-95% against *V. cholerae* and 93-94% against *V. paracholerae*. Since the results are close to the cut-off value of 96% ANI for two genomes to belong to the same species [22], we complemented our ANI results with dDDH. The GGDC package was used to calculate percent dDDH *in silico* to mimic wet lab-based DDH [23]. Within *V. tarriae* sp. nov., the dDDH values were 81.5-85%, whereas values were 60.6-67.9% and 59.8-67% when compared with *V. cholerae* and *V. paracholerae*, respectively (Fig. 1).

**Figure 1:**
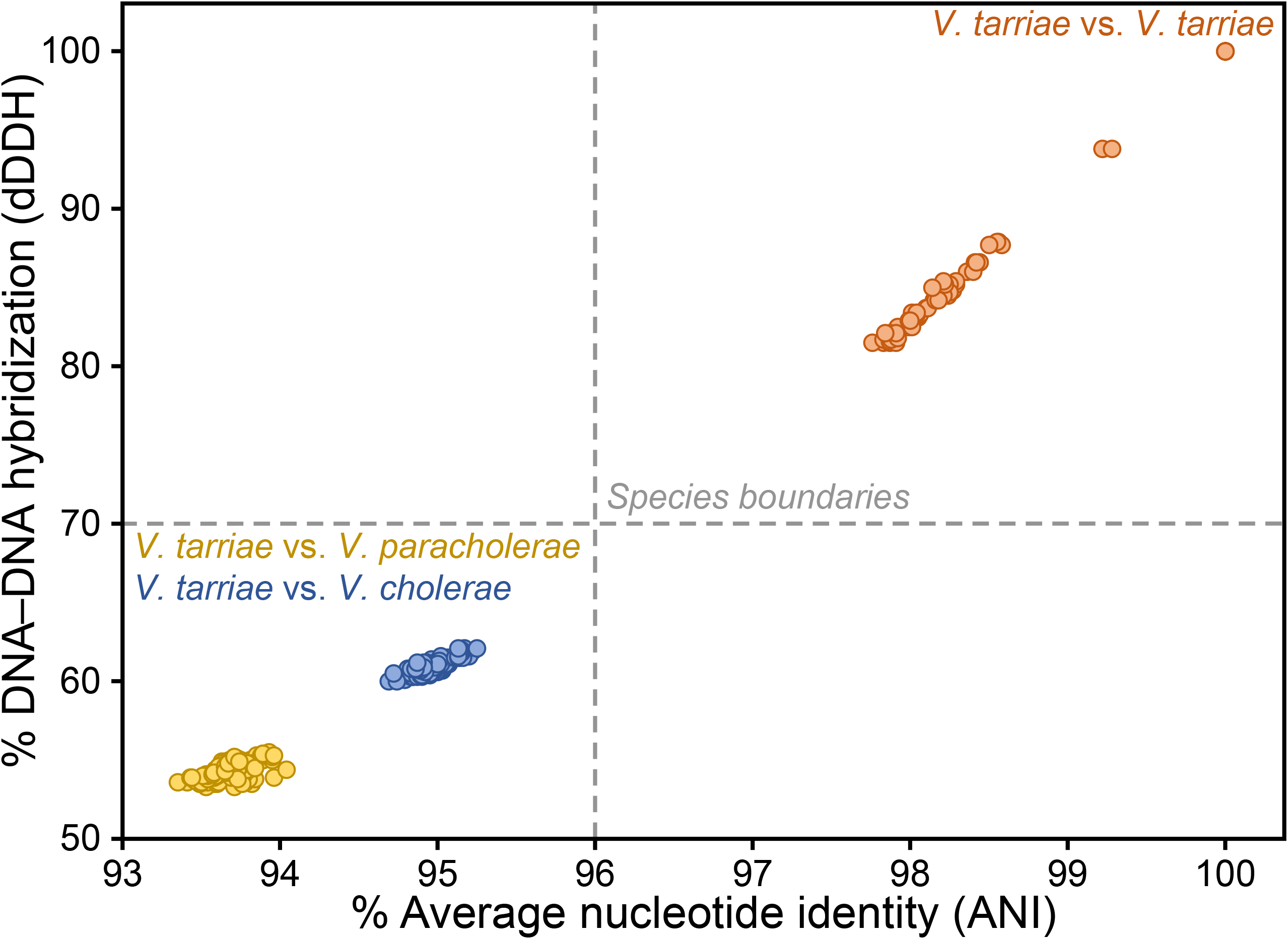
Whole-genome comparisons among isolates of *V. tarriae* sp. nov., *V. cholerae* and *V. paracholerae*. Pairwise percentage value of digital DNA-DNA hybridization(dDDH) and avergae nucleotide identity (ANI) comparisons between and group and within groups are shown. Genomes of *V. tarriae* sp. nov. falling below the species boundary cut-offs indicating that they are a distinct species.

MLSA further supports our proposal for defining a novel species. Single-copy, protein-coding core genes are used as alternatives to 16S rRNA gene sequences for the identification and phylogenetic analysis of various species of the genus *Vibrio*, since there is a lack of species-level resolution using 16S rRNA gene sequence [22, 28]. The *V. tarriae* sp. nov. isolates form a monophyletic clade that is distinct from *V. cholerae* and other *Vibrio* species, with 100% bootstrap support using six housekeeping genes (*adk, gyrB, pyrH, pgi, recA* and *rpoA)* (Fig. 2A). These genes have been used for the taxonomic characterization of vibrios and to describe novel species within the genus [8, 20, 28]. The average patristic distance calculated from this tree between *V. tarriae* sp. nov. isolates and *V. cholerae* and *V. paracholerae* isolates is 0.06 and 0.09, respectively, while lower average distances of 0.03 are obtained when comparing isolates within the species.

**Figure 2:**
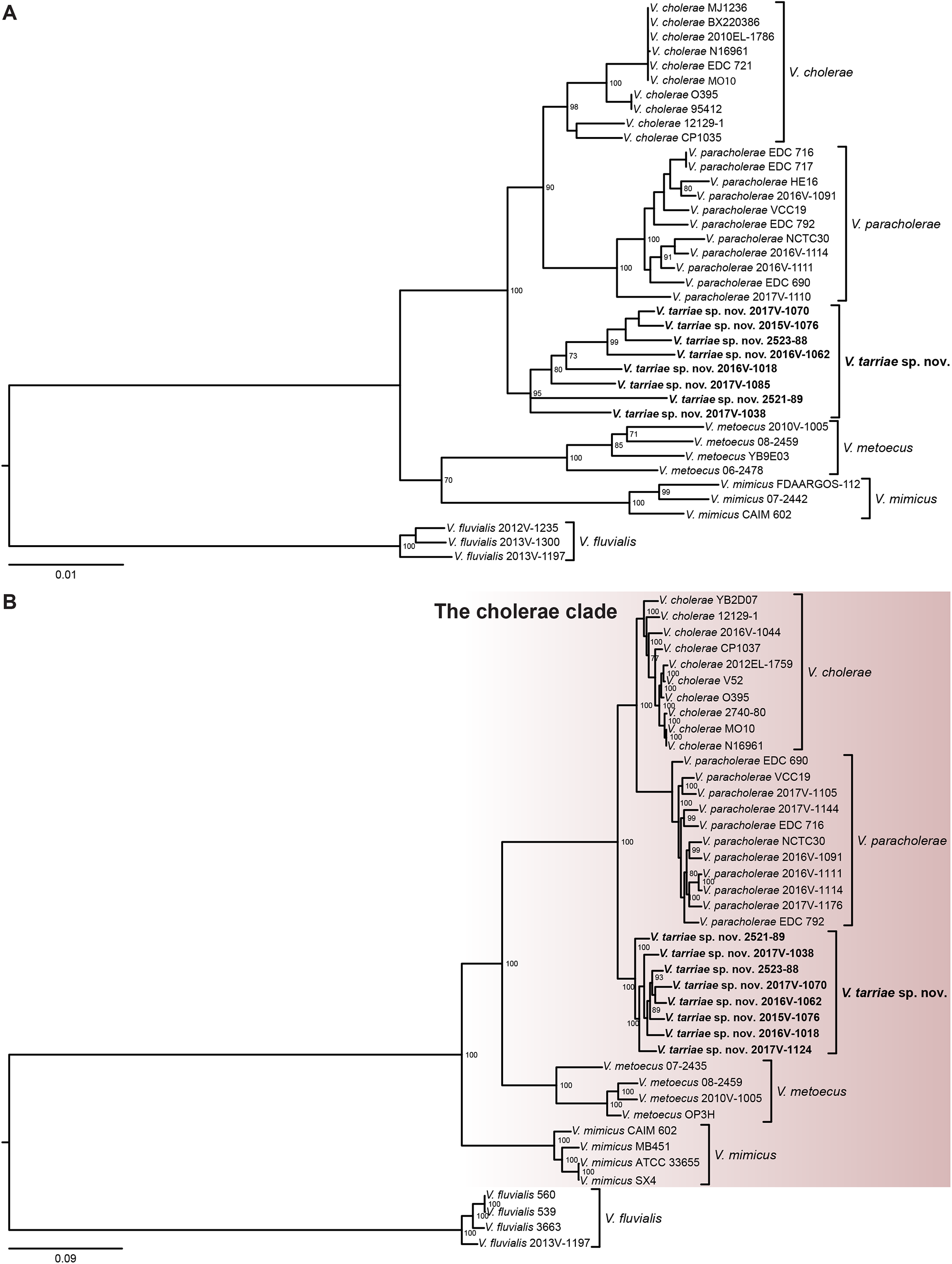
Phylogenetic relationship of *Vibrio tarriae* sp. nov. and its closest relatives. A) Multi locus sequence analysis (MLSA) performed based on the concatenated alignment of DNA sequences of six protein coding housekeeping genes: *adk, gyrB, pyrH, pgi, recA* and *rpoA* were used for the analysis. Bootstrap values >70% are shown in the nodes of relevant branches. The scale bar represents nucleotide substitutions per site. B) The maximum likelihood phylogenetic tree was constructed from the core genome alignment of ≈ 2.1M bp using GTR gamma substitution model. The Cholerae clade was marked by the shaded box. Bootstrap values >70% are shown in the nodes of relevant branches. The scale bar represents nucleotide substitutions per site.

To further demonstrate this distinction, a phylogeny was reconstructed based on the core genome sequence of these three species (Fig. 2B). Core genome phylogeny shows the distinct monophyletic clustering of *V. tarriae* sp. nov. isolates from other *Vibrio* species. This placedbthe *V. tarriae* sp. nov. lineage into the context of a larger *Vibrio* phylogeny, showing that the novel species is distinct from all *Vibrionaceae* that have been characterized to date but fall into the Cholerae clade. It is more closely related to *V. cholerae* than *V. metoecus* or *V. mimicus* but forms an outgroup to the *V. cholerae* and *V. paracholerae* sister species (Fig. 2B). This distinction is further confirmed by ANI and percent DDH below 96% and 70%, respectively, between *V. tarriae* sp. nov. and other closely related species (Fig. 1).

### Genetic basis for the divergence and ecology of the novel species

Interestingly, there were some signatures of divergence in terms of the gene content in *V. tarriae* sp. nov. from all other species in the cholerae clade (*V. cholerae, V. paracholerae, V. metoecus* and *V. mimicus)*. While genome phylogeny confirms the close relatedness of *V. tarriae* sp. nov. to *V. cholerae* and *V. paracholerae*, gene content analysis revealed a few species-specific genetic traits which differentiates it from the two closest species (Table 3). There were at least eight gene clusters which were present in all nine *V. tarriae* sp. nov. strains but absent in all *V. cholerae* and *V. paracholerae* strains tested (Fig. 3). Notably, the siderophore system had differences in *V. tarriae* sp. nov. compared with the other two most closely related species. Even though *V. tarriae* sp. nov. forms a monophyletic clade with *V. cholerae* and *V. paracholerae* (Table 3 and Fig. 2), gene content of this siderophore system is more closely related to *V. metoecus* and *V. mimicus*. Specifically, a gene cluster (GC-1) containing genes (CEQ48_09230-CEQ48_09275) for aerobactin siderophore common to *V. metoecus* and *V. mimicus* are present in *V. tarriae* sp. nov. but absent in *V. cholerae* and *V. paracholerae* (Fig. 3 and Table 3). On the other hand, *V. tarriae* sp. nov. strains lack genes for the vibriobactin siderophore system present in *V. cholerae* and *V. paracholerae* (Table 3).

**Table 3.**
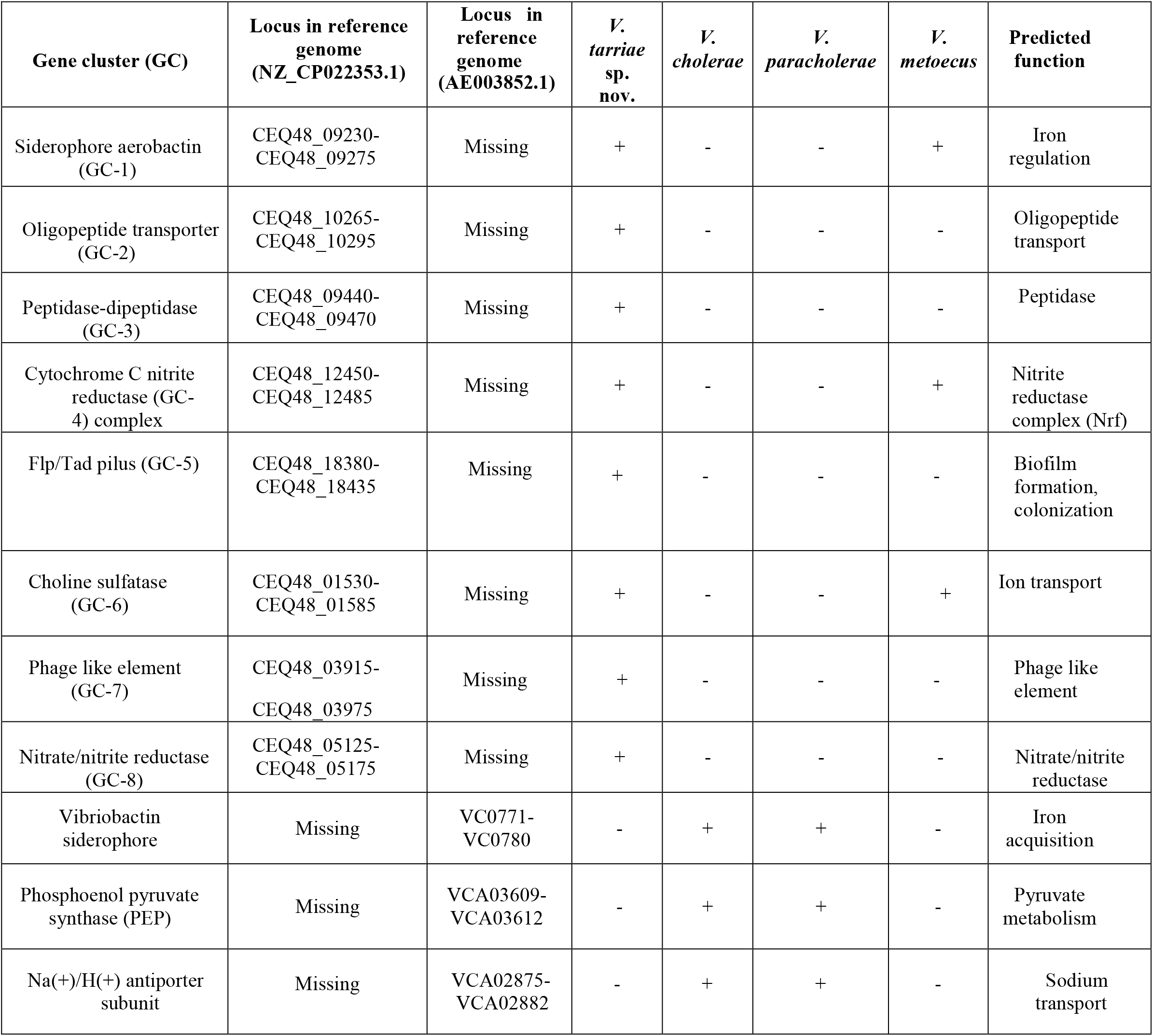
Major genetic traits differentiating *Vibrio tarriae* sp. nov. from its closest relatives: *Vibrio cholerae, Vibrio paracholerae and Vibrio metoecus*. Reference genomes N16961 (*V. cholerae*) and 2521-89 (*V. tarriae* sp. nov.) were used for determining locus positions of the gene clusters.

**Figure 3:**
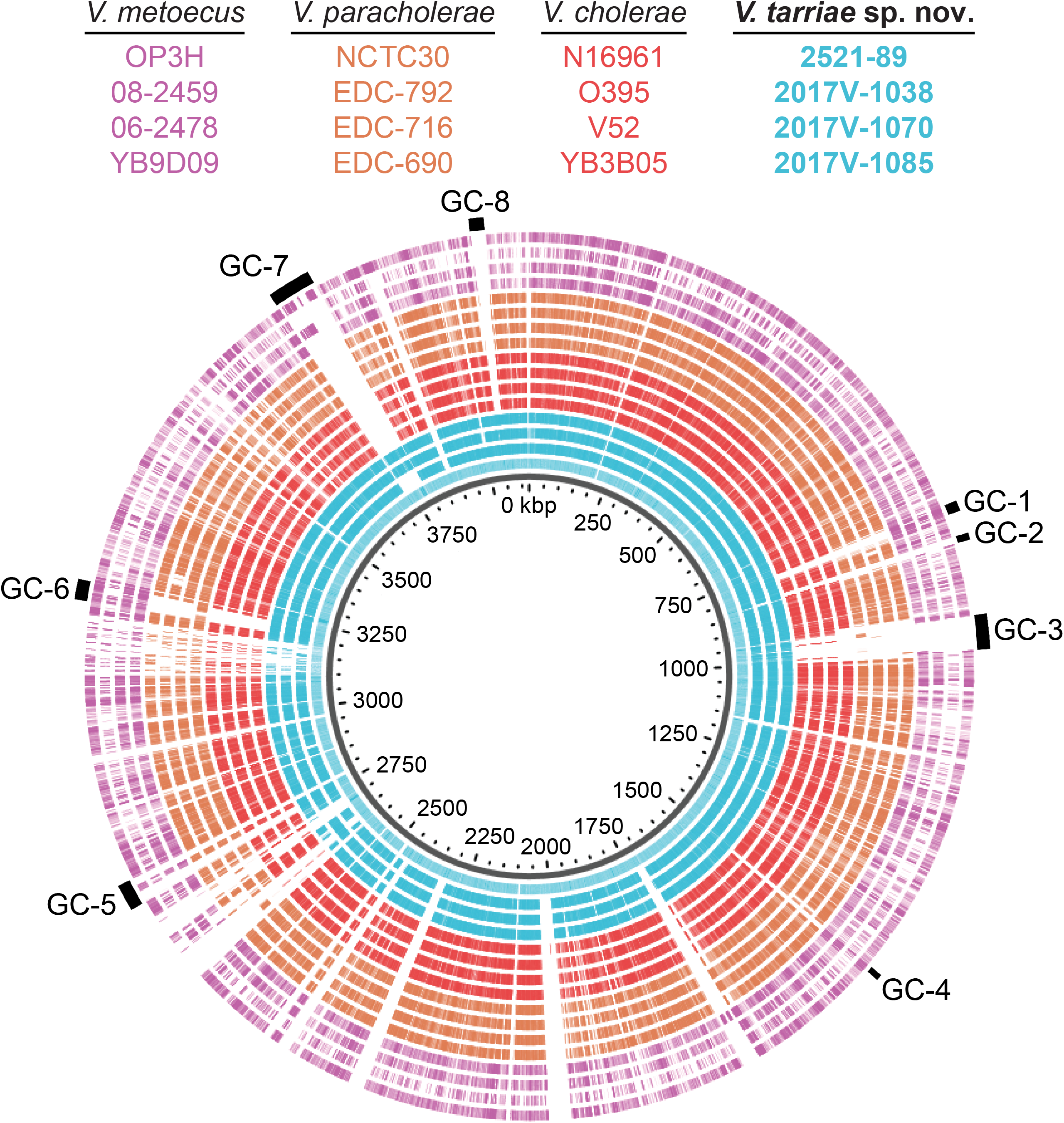
BLAST atlas of *V. tarriae* sp. nov. along with *V. cholerae, V. paracholerae* and *V. metoecus* genomes. *V. tarriae* sp. nov. strain 2521-89 was used as the reference sequence for BLASTN comparisons. Each colored ring represents a genome. Outermost black bars indicate the gene clusters (GC’s) present in *V. tarriae* sp. nov. which are absent in any other sister species (Table 3). GC-1, Gene cluster-1; GC-2, Gene cluster-2; GC-3, Gene cluster-3; GC-4, Gene cluster-4; GC-5, Gene cluster-5; GC-6, Gene cluster-6; GC-7, Gene cluster-7; GC-8, Gene cluster-8.

Like the siderophore system, two additional gene clusters (GC-4 and GC-6) were present in *V. tarriae* sp. nov., *V. metoecus* but absent in most *V. cholerae* and *V. paracholerae*. Gene cluster GC-6 (CEQ48_01530-CEQ48_01585) including genes for Choline-sulfatase (EC 3.1.6.6) and Lacto-N-biose phosphorylase (EC 2.4.1.211) were present in all the *V. tarriae* sp. nov. isolates but absent in *V. cholerae* (n=22) and most (21 out of 22) of the *V. paracholerae* isolates (Fig. 3 and Table 3). The other gene cluster (GC-4) common to the *V. tarriae* sp. nov. isolates contained genes (CEQ48_12450-CEQ48_12485) for a nitrate reductase complex (Nrf) that can be important for the adaptation of the bacterium in hypoxic conditions like the human intestine [31].

Interestingly, there was an additional nitrate reductase complex gene cluster (GC-8) not found in any of the sister species within the cholerae clade but present in all the *V. tarriae* sp. nov. isolates. Including GC-8, five gene clusters (GC-2, GC-3, GC-5, GC-7 and GC-8) found in all *V. tarriae* sp. nov. strains are not common in any of its close relatives: *V. cholerae, V. paracholerae, V. metoecus* and *V. mimicus* (Table 3). GC-2 and GC-3 are most likely to be involved in peptide catabolism as GC-2 (CEQ48_10265-CEQ48_10295) contains genes for peptide transport system and GC-3 (CEQ48_09440-CEQ48_09470) contain genes for peptidases, respectively (Fig. 3 and Table 3). GC-5, a gene cluster (CEQ48_18380-CEQ48_18435) encoding genes for Tad pilus, is also an indicator of divergence found only in *V. tarriae* sp. nov. and absent in other species within the clade (Fig. 3 and Table 3). Tad pili are associated with biofilm formation and surface colonization in several other bacteria including the human pathogen *Vibrio vulnificus* [32]. GC-7 (CEQ48_03915-CEQ48_03975) encodes a putative prophage of ∼10 kbp. GC-8 (CEQ48_05125-CEQ48_5175) is a ∼15 kbp region containing genes for the nitrate reductase complex, to which the closest homolog was found in opportunistic human pathogen *Vibrio navarrensis*.

These divergences in the gene content including siderophore, nitrate reductase, nutrient transporter, attachment pili and putative prophage-related gene repertoires can be important mediators for the diversification of substrate utilization, ecological interactions, and association with animal host among the closely related species of the cholerae clade. For example, nitrite reductase have been thought to be required for growth in anoxic conditions (i.e., animal host intestine, using nitrate reduction) [31]. Production of unique siderophores can provide competitive advantage to a species from its neighbors in a shared habitat [33]. Also, possession of additional varieties of siderophore receptors can be useful in scavenging iron captured by siderophores secreted by other bacteria in an iron limiting conditions such as the human intestine [34]. Other than the vibriobactin siderophore, two other gene clusters potentially related to pyruvate metabolism and sodium transport (Table 3) commonly found in *V. cholerae* and *V. paracholerae* were missing in *V. tarriae* strains. Seven out of nine isolates of *V. tarriae* sp. nov. analyzed in this study were isolated from human clinical samples (Table 1), which suggests that the species might be an opportunistic pathogen to humans. While the pathogenesis and prognosis of the disease could not be clearly defined, genetic similarities and differences observed in *V. tarriae* sp. nov. to and from its well-known close relative *V. cholerae* and other ancestrally related species within the cholerae clade could be important clues in understanding virulence and pathogenesis mechanisms.

## Conclusion

The Cholerae clade is an important group of bacteria containing the well-known environmental pathogen *V. cholerae* and several potentially human-associated *Vibrio* species. Exact phylogenetic structuring and genetic signatures of the clade is being refined with the discovery and description of novel closely related species potentially involved in human infections. The inclusion of the *V. tarriae* sp. nov. in the group updates our understanding of the diversity of the clade as well as the genus *Vibrio*. Phenotypic and genetic differences observed in *V. tarriae* s. nov. make it an interesting model for studying the evolution and diversification of *Vibrionaceae* in general. So far, *V. tarriae* sp. nov. was isolated from environmental and human samples from the U.S. only; exploration of more genomic and demographic data would shed light on whether it is endemic to the U.S. or distributed globally like several species of the Cholerae clade.

### Description of *V. tarriae* sp. nov

*Vibrio tarriae* (tarr’i.ae. N.L. gen. fem. n. tarriae of tarr., named after Dr. Cheryl Tarr, in recognition of her contribution to the study of *Vibrio* diversity and exploration of atypical vibrios).

Cells are Gram-negative, curved, motile rods, 0.64–0.78×1.48–1.68 µm in size, which produce convex, smooth, circular, entire, cream colonies on TSB agar and yellow sucrose fermenting *V. cholerae* like colonies on TCBS agar. Growth is observed in TSB at 30 °C with salt concentrations in the range of 0–6.5 % NaCl; no growth occurs in the presence of 10 % NaCl. Growth is also observed in TSB with 1.5 % total NaCl concentration at a temperature range of 30–40 °C, and no growth occurs at 4 °C and 45 °C. The ability to utilize D-lactose, *N*-acetyl-D-glucosamine and pectin as well as the lack of ability to utilize D-mannose, D-serine, citric acid, D-ribose, propionic acid, D-alanin, mono-methyl succinate and caproic acid as the sole carbon and energy source differentiates *V. tarriae* from *V. cholerae*. On the other hand, *V. tarriae* can be differentiated from *V. paracholerae* by the ability to utilize N-acetyl-D-glucosamine and inability to utilize α-cyclodextrin as the sole carbon and energy source.

*V. tarriae* can produce indole and β-glucosidase as well as reduce nitrate to nitrite, positive for glucose fermentation, lysine decarboxylase and ornithine decarboxylase. This strain produces acid from the fermentation of mannitol but not arabinose, and is arginine dihydrolase-and urease-negative. On TCBS agar, formation of *V. cholerae*-like yellow circular colonies is observed, while flat, smooth, circular colonies of creamy-white color on TSB agar is observed. This strain is also positive for carbon utilization from α-D-glucose, β-galactose, citrate, D-fructose, D-glucoronic acid, D-maltose, D-trehalose, dextrin, gelatin, N-acetyl-D-galactosamine, sucrose, D-lactose and pectin. Diversity between strains exists for utilization of L-asparagine, L-glutamine, glycyl-L-aspartic acid, N-acetyl-D-Glucosamine, glycerol, D, L-malic acid, D-glucose-1-phosphate, L-serine, L-threonine, succinic scid, D-glucuronic scid, Tween 40, D-psicose, D-galactose, D-gluconic acid, α-keto-glutaric acid, α-hydroxy glutaric acid-γ-lactone, bromo succinic acid, L-alanyl-glycine, L-aspartic acid, D, L-α-glycerol-phosphate, L-proline, β-methyl-D-glucoside, N-acetyl-β-D-mannosamine, L-galactonic acid-γ-lactone, methyl pyruvate, D-mannitol, L-glutamic acid, thymidine, gentiobiose, raffinose, capric acid, salicin, D-glucosamine and laminarin.

The type strain 2521-89^T^ (= DSM 112461^T^ = CCUG 75318^T^) was isolated from a lake in New Mexico, U.S. Based on whole-genome sequencing, the G+C content of the type strain is 47.1%.

## Conflict of interests

The authors declare no conflicts of interest.

## Data availability

All genomic data is available in public databases and can be accessed using the Genbank accession numbers listed in Table 1 and available on request.

## Funding information

M. T. I. and K.L were supported by Alberta Innovates Technology Futures. Y. B. was supported by the Natural Sciences and Engineering Research Council of Canada. Y. B. was also supported by the Canadian Institute for Advanced Research.

## Notes

### Competing Interest Statement

The authors have declared no competing interest.

